## Supplemental Figures for "Differential modularity of the mammalian *Engrailed 1* enhancer network directs eccrine sweat gland development"

**This PDF file includes:**

Figures S1 to S4
Table S1
SI References

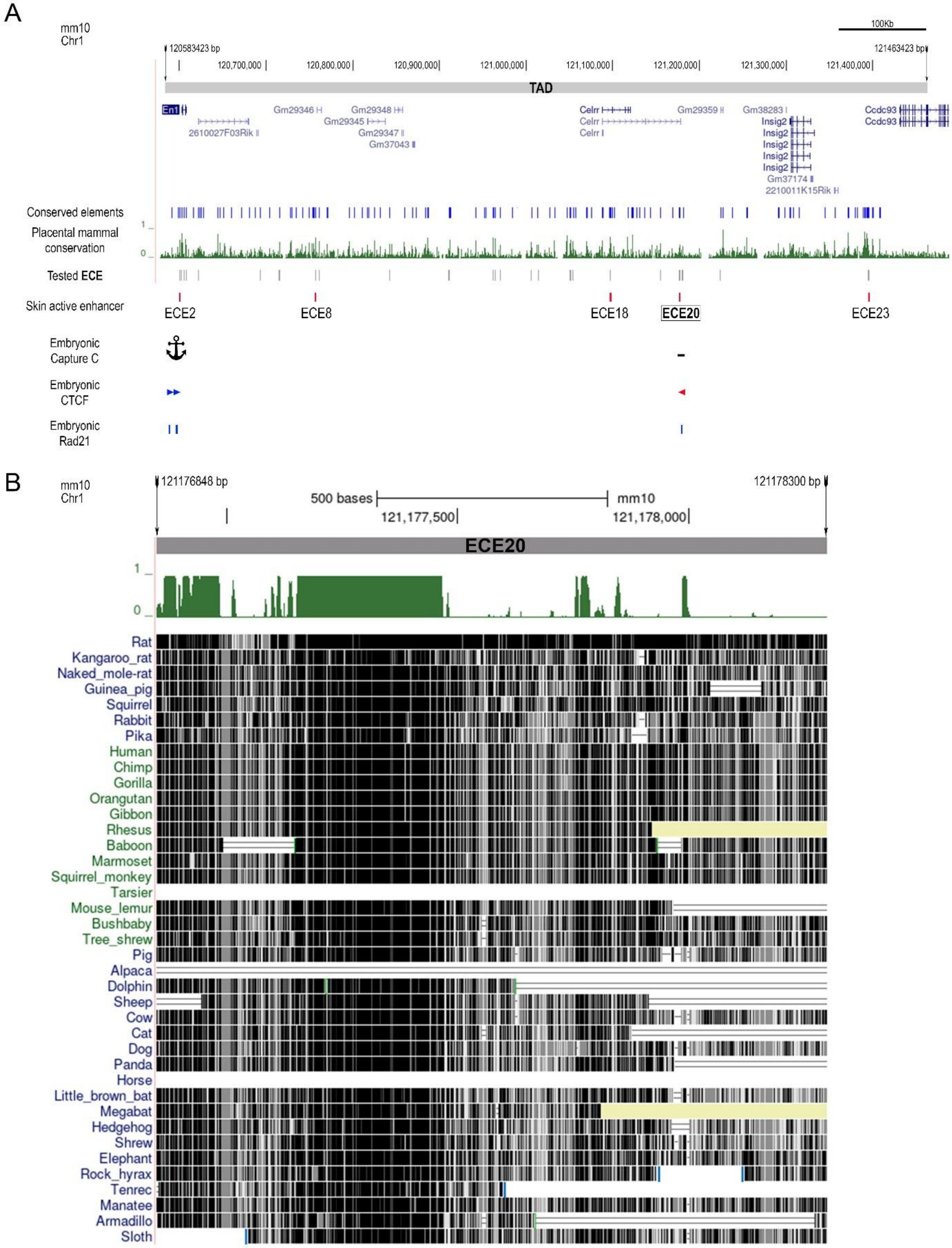
A
mm10
Chr1
100Kb
120583423 bp
121463423 bp
120,700,000
120,800,000
120,900,000
121,000,000
121,100,000
121,200,000
121,300,000
121,400,000
TAD
En1
2610027F03Rik
Gm29346
Gm29348
Gm29345
Gm29347
Gm37043
Celrr
Celrr
Celrr
Gm29359
Gm38283
Insig2
Insig2
Insig2
Insig2
Insig2
Gm37174
2210011K15Rik
Ccdc93
Ccdc93
Conserved elements
Placental mammal conservation
1
0
Tested ECE
Skin active enhancer
ECE2
ECE8
ECE18
ECE20
ECE23
Embryonic Capture C
Embryonic CTCF
Embryonic Rad21
B
mm10
Chr1
121176848 bp
121178300 bp
500 bases
mm10
121,177,500
121,178,000
ECE20
1
0
Rat
Kangaroo_rat
Naked_mole-rat
Guinea_pig
Squirrel
Rabbit
Pika
Human
Chimp
Gorilla
Orangutan
Gibbon
Rhesus
Baboon
Marmoset
Squirrel_monkey
Tarsier
Mouse_lemur
Bushbaby
Tree_shrew
Pig
Alpaca
Dolphin
Sheep
Cow
Cat
Dog
Panda
Horse
Little_brown_bat
Megabat
Hedgehog
Shrew
Elephant
Rock_hyrax
Tenrec
Manatee
Armadillo
Sloth

C

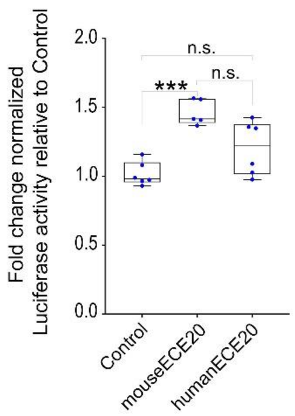


**Fig. S1. Genomic location of the Engrailed 1 Candidate Enhancer (ECE) actives during skin development and characteristics of the ECE20 element. (A)** Positions of the 209 conserved elements (blue lines) within the En1 TAD previously identified by phastCons, a software which detects string of high conservation given an alignment of species (in this case placental mammals). PhastCons score graph is depicted in green. Positions of the 23 ECEs (grey lines) tested *in vivo*, and the 5 ECEs with ectodermal activity in mice. Positive ECEs were called based on detection of enhancer activity in *En1* expressing cells of the volar paw skin of transgenic mice[1]. Genomic location corresponding to Capture-C, CTCF and RAD21 peaks from developing limbs and midbrain mice identified in Andrey et al. [2]. The viewpoint located at the En1-promoter (anchor) was used to generate the Capture-C spatial map interaction [2]. Dash line represent the position of the only peak of interaction with the En1-promoter. Blue and red arrows indicate direction of CTCF peaks. Blue lines indicates RAD21 peaks [2]. **(B)** UCSC genome browser screenshot illustrating the alignment of placental mammal sequences centered on ECE20. mm10 was use as genome base. Placental conservation by means of PhasCons is depicted. **(C)** Comparative quantitative activity of mouse and human ECE20 orthologs in cultured human GMA24F1A keratinocytes, an immortalized human skin cell line that endogenously expressed *EN1*. Fold change normalized luciferase activity relative to Control (empty vector) is plotted. In **(C)** significance assessed by one-way *ANOVA* and Tukey-adjusted P-values are reported. ***P<0.001, n.s. not significant.

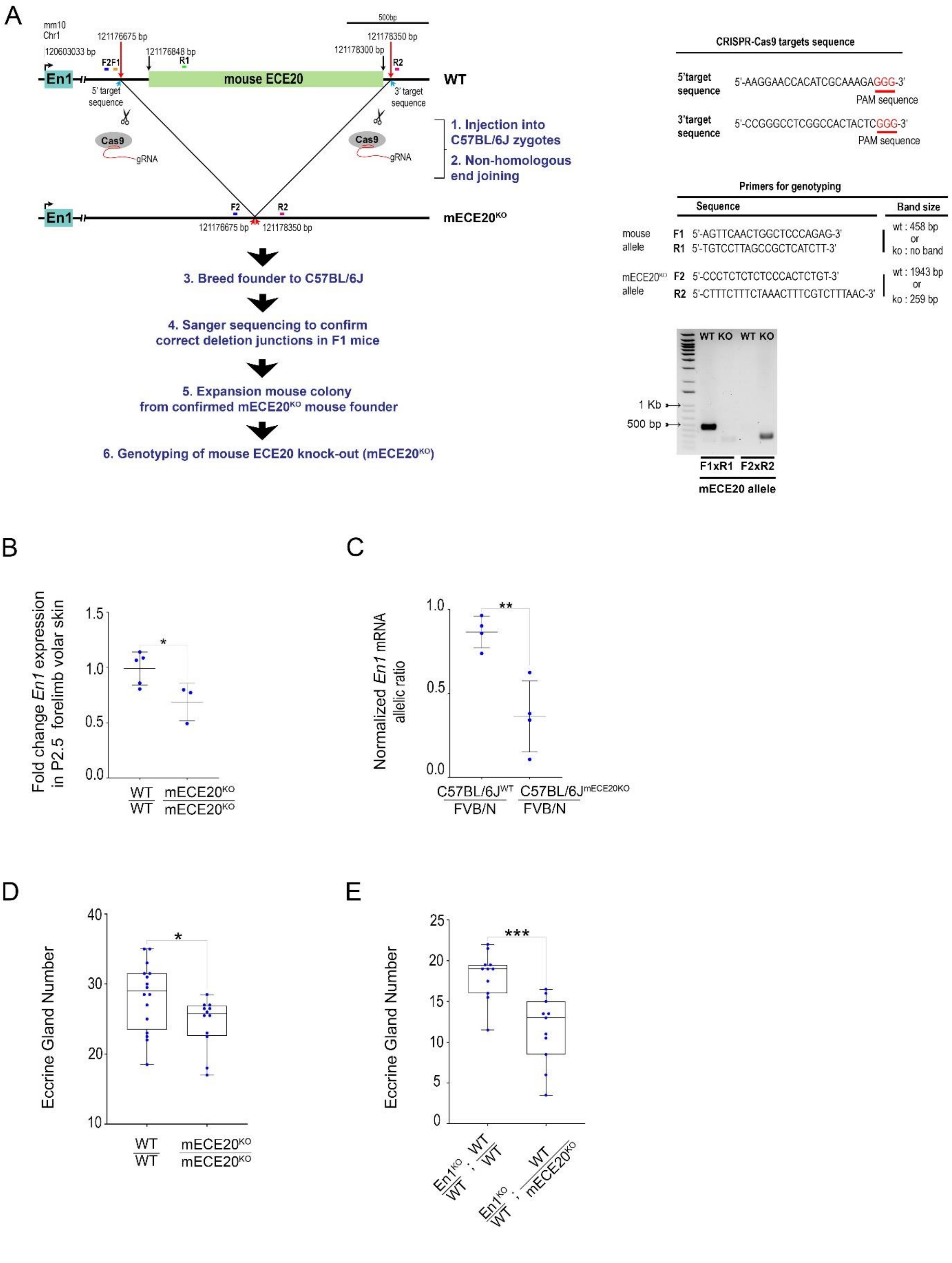

A
mm10
Chr1
120603033 bp
121176675 bp
121176848 bp
121178300 bp
121178350 bp
500bp
F2
F1
R1
R2
En1
mouse ECE20
WT
5' target sequence
3' target sequence
Cas9
gRNA
1. Injection into C57BL/6J zygotes
2. Non-homologous end joining
mECE20KO
3. Breed founder to C57BL/6J
4. Sanger sequencing to confirm correct deletion junctions in F1 mice
5. Expansion mouse colony from confirmed mECE20KO mouse founder
6. Genotyping of mouse ECE20 knock-out (mECE20KO)
CRISPR-Cas9 targets sequence
5'target sequence 5'-AAGGAACCACATCGCAAAGAGGG-3'
PAM sequence
3'target sequence 5'-CCGGGCCTCGGCCACTACTCGGG-3'
PAM sequence
Primers for genotyping
Sequence
Band size
mouse allele
F1 5'-AGTTCAACTGGCTCCCAGAG-3'
R1 5'-TGTCCTTAGCCGCTCATCTT-3'
wt : 458 bp or ko : no band
mECE20KO allele
F2 5'-CCCTCTCTCTCCCACTCTGT-3'
R2 5'-CTTTCTTTCTAAACTTTCGTCTTTAAC-3'
wt : 1943 bp or ko : 259 bp
WT KO WT KO
1 Kb
500 bp
F1xR1
F2xR2
mECE20 allele
B
Fold change En1 expression in P2.5 forelimb volar skin
1.5
1.0
0.5
0.0
*
WT/WT
mECE20KO/mECE20KO
C
Normalized En1 mRNA allelic ratio
1.0
0.5
0.0
**
C57BL/6JWT/FVB/N
C57BL/6JmECE20KO/FVB/N
D
Eccrine Gland Number
40
30
20
10
*
WT/WT
mECE20KO/mECE20KO
E
Eccrine Gland Number
25
20
15
10
5
0
***
En1KO/WT ; WT/WT
En1KO/WT ; WT/mECE20KO

**Fig. S2. Generation ECE20 knock-out mouse and characterization of *En1* expression and eccrine gland phenotypes in the volar forelimb. (A)** Generation of an ECE20 knock-out mouse (mECE20$^{KO}$) by CRISPR-Cas9 mediated genome editing. CRISPR-Cas9 target sequence and genotyping strategy are shown. Correct deletion junctions were confirmed by Sanger sequencing of F1 pups. **(B)** Fold change in *En1* mRNA by qRT-PCR in P2.5 volar forelimb skin of wildtype (WT / WT), and mECE20$^{KO}$ homozygote (mECE20$^{KO}$/ mECE20$^{KO}$) mice relative to wildtype. **(C)** Normalized *En1* mRNA allelic ratio in volar forelimb of wildtype at P2.5 of wildtype (C57BL/6J$^{WT}$ / FVB/N) and mECE20$^{KO}$ (C57BL/6J(mECE20$^{KO}$) / FVB/N) hybrid mice. Ratios were normalized to the allelic ratio in F1 genomic DNA. Each point represents the mean value across three technical replicates of biological samples consisting of pooled P2.5 volar skins from both forelimbs of two or three mice. **(D)** Quantification of IFP eccrine glands in WT / WT and mECE20$^{KO}$/ mECE20$^{KO}$ mice. **(E)** Quantification of IFP eccrine glands in En1 $^{KO}$ / WT; WT / WT and En1 $^{KO}$ / WT ; WT / mECE20$^{KO}$ mice. In **(B, C)** dots represent an individual biological replicate. In **(D, E)** each point represents the average number of eccrine glands in the IFP across both forelimbs of a mouse. In **(B, C)** mean (line) with standard deviation are plotted. In **(D, E)** the median (line) and maximum and minimum are reported for each genotype. In **(B-E)** significance assessed by a two-tailed T-test. ***P<0.001, ** *P*<0.01, * *P*<0.05. (KO) knock-out. In **(B, C)** *Rlp13a* was used as housekeeping transcript for normalization.

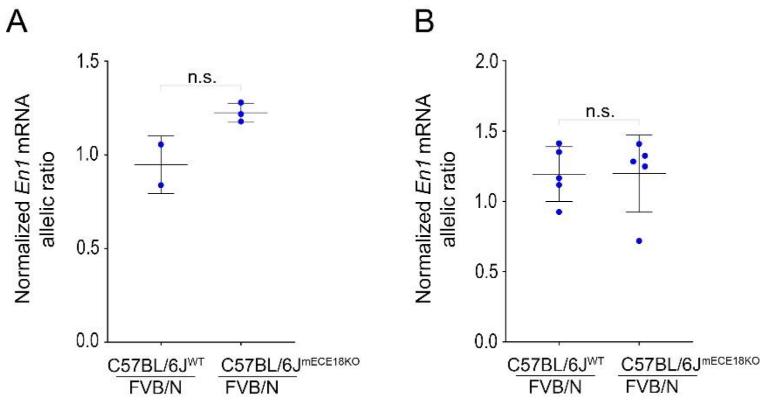


**Fig. S3. Effect of mECE18 knock-out on En1 expression in the mouse embryonic midbrain-hindbrain and limb-bud.** Normalized *En1* mRNA allelic ratios in forelimb autopods **(A)** and midbrain-hindbrain **(B)** of wildtype (C57BL/6J$^{WT}$ / FVB/N) and mECE18$^{KO}$ (C57BL/6J$^{mECE18KO}$ / FVB/N) hybrid mice are plotted. Ratios normalized to genomic DNA allelic ratio. Each point represents the mean value across three technical replicates of a pooled of three or four mice for embryonic limb-bud **(A)** or individual dissections of midbrain-hindbrain **(B)** at E10.5. In **(A, C)** mean (line) with standard deviation is reported. In **(A, B)** significance assessed by a two-tailed T-test. n.s. not significant. (KO) knock-out.

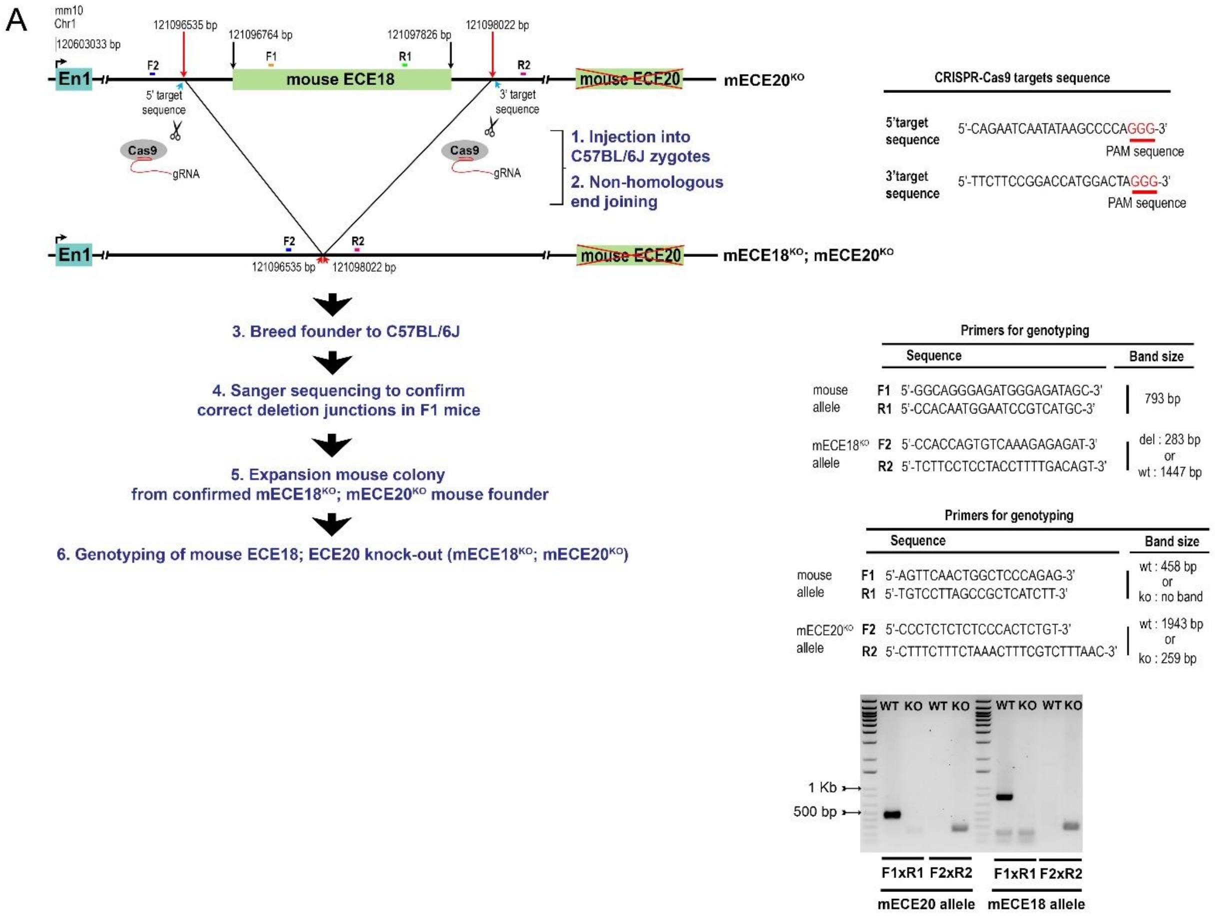


**Fig. S4. Generation of ECE18; ECE20 compound knock-out mice. (A)** Generation of an ECE18; ECE20 compound knock-out mouse (mECE18$^{KO}$; mECE20$^{KO}$) by CRISPR-Cas9 mediated genome editing. CRISPR-Cas9 target sequence and genotyping strategy are shown. Correct deletion junctions were confirmed by Sanger sequencing of F1 pups.

### References

1. Aldea D, Atsuta Y, Kokalari B, Schaffner SF, Prasasya RD, Aharoni A, et al. Repeated mutation of a developmental enhancer contributed to human thermoregulatory evolution. Proc Natl Acad Sci U S A. 2021;118: e2021722118. doi:10.1073/pnas.2021722118

2. Andrey G, Schöpflin R, Jerković I, Heinrich V, Ibrahim DM, Paliou C, et al. Characterization of hundreds of regulatory landscapes in developing limbs reveals two regimes of chromatin folding. Genome Res. 2017;27: 223–233. doi:10.1101/gr.213066.116

**Table S1:**

Primers used to subclone ECE20 orthologs in mouse transgenic assays[#]

| ECE | Forward sequence | Reverse sequence |
|---|---|---|
| mouse-ECE20 | accaattgctcgaggCACATTCAAGGTCAATG | gtcaagcttccattatatagGCAGCAGTGAGTGTG |
| human-ECE20 | agtcgaccaattgctcgaggTACATCCAAGGCCAGTTTCTCC | cggtcaagcttccattatatagGCTACCGTGGGCGCCTGA |

[#]Lower case sequence indicates homology arms to Stagia3 vector

Primers used to subclone ECE20 orthologs into bidirectional luciferase reporter vector [##]

| ECE | Forward sequence | Reverse sequence |
|---|---|---|
| mouse-ECE20 | agagatttagaatgacaggcCACATTCAAGGTCAATGTCTCC | cttccattatatagaattCCGCAGCAGTGAGTGTGCGCGC |
| human-ECE20 | agagatttagaatgacaggcTACATCCAAGGCCAGTTTCTCCAA | aagcttccattatatagaattCCGCTACCGTGGGCGCCTGAGCAGAGCC |

[##]Lowercase sequence indicates homology arm to vector

Primers used for qRT-PCR primer sequences

| Name | Specie | Forward sequence | Reverse sequence |
|---|---|---|---|
| En1 | Mouse | GTGGTCAAGACTGACTCACAGC | GCTTGTCTTCCTTCTCGTTCTT |
| Rpl13a | Mouse | CAGTGCGCCAGAAAATGC | GAAGGCATCAACATTTCTGGAA |

Primers used for allelic discrimination assay

| Name | Specie | Forward sequence | Reverse sequence |
|---|---|---|---|
| En1 | Mouse | GAGCAGCTGCAGAGACTCAA | CTCGCTCTCGTCTTTGTCCT |
